# The Intestinal Clock Regulates Host Metabolism through the Fiber-Dependent Microbiome and Macronutrient Transcriptome

**DOI:** 10.1101/2023.04.11.534733

**Authors:** Marjolein Heddes, Yunhui Niu, Baraa Altaha, Karin Kleigrewe, Chen Meng, Dirk Haller, Silke Kiessling

## Abstract

Circadian disruption, e.g. through shift work, causes microbial dysbiosis and increases the risk of metabolic diseases. Microbial rhythmicity in mice depends on a functional intestinal clock and frequent jetlag as well as high-caloric energy intake induces loss of these oscillations. Similarly, arrhythmic microbiota was found in obese and T2D populations. However, the interplay between the intestinal circadian clock, the microbiome, diet and host metabolism is poorly understood.

In intestinal-specific *Bmal1* knockout mice (*Bmal1^IEC-/-^*) we demonstrate the relevance of the intestinal clock in microbiome oscillations and host and microbial nutrient metabolism. Microbiota transfer from *Bmal1^IEC-/-^* mice into germ-free recipients led to obesity, reflected by increased bodyweight and fat mass. Western diet-fed *Bmal1^IEC-/-^ mice* increased bodyweight likely through mechanisms involving the intestinal clock-control of lipid and hexose transporters. Additionally, we identified dietary fiber as novel link between circadian microbial rhythmicity, intestinal clock functioning and host physiology. Thus, revealing the potential of fiber-rich diet intervention as a non-invasive strategy targeting microbial oscillations in metabolic disease prevention.

## Introduction

Circadian (∼24h) clocks play an important role in adjusting behavior and physiology to recurring environmental changes (e.g. food availability). Besides the central clock located in the hypothalamus, most tissues and even single cells harbor circadian oscillators, so called peripheral clocks (Schibler, 2003). On the molecular level the circadian clock consist of core clock genes forming an autoregulatory transcriptional-translational feedback loop, thereby driving circadian oscillations of genes involved in various pathways (Takahashi, 2017). While the central clock in the hypothalamus is mainly entrained by light, peripheral clocks, including pancreas, liver, adipose tissue and the gastrointestinal (GI) tract, are predominantly entrained by the timing and composition of nutrient intake (Damiola et al., 2000; Desmet et al., 2021; Pickel and Sung, 2020). These metabolic relevant organs are responsible for digestion, absorption as well as nutrient utilization in a circadian manner (Pickel and Sung, 2020). Particularly, the GI tract plays an essential role in circadian metabolic homeostasis, since it is the first organ encountering nutrients (Hoogerwerf et al., 2008; Sladek et al., 2007). Digestion and absorption of major nutrients, such as fat, carbohydrates and proteins within the GI system are clock controlled through the rhythmic expression of ion, sugar, peptides, triglyceride and cholesterol transporter genes (Balakrishnan et al., 2012; Iwashina et al., 2011; Pacha and Sumova, 2013; Pan and Hussain, 2007; 2009; Pan et al., 2004). In human cohorts, global circadian disruption e.g. through shift work, results in altered feeding patterns and correlated to metabolic diseases. Moreover, another important link between metabolism and diet is the microbiome. In mouse models system-wide and tissue-specific genetic loss of *Bmal1,* a major circadian clock gene, results in alteration in glucose and lipid metabolism, adipocyte differentiation and arrhythmic microbial metabolite production, highlighting the importance of a functional circadian clock in host metabolism (Heddes et al., 2022; Hodge et al., 2015; Onuma et al., 2022; Segers et al., 2020; Shimba et al., 2005; Yu et al., 2021). Recently we showed in intestinal epithelial cell (IEC)-specific *Bmal1*-deficient mice that the intestinal clock plays an important role in driving circadian microbiota composition and function e.g. by regulation of microbial-associated molecular patterns (MAMPs) (Heddes *et al*., 2022). MAMPs, including among others short-chain fatty acids (SCFAs), amino acids as well as bile acids (BAs), signal to the host’s epithelium thereby regulating intestinal clock gene expression involved in lipid and glucose uptake and metabolism (Frazier et al., 2020; Govindarajan et al., 2016; Tahara et al., 2018). These results suggest the importance of the intestinal clock in microbiome mediated metabolic homeostasis. This hypothesis is strengthened by our transfer experiments suggesting that loss of microbial rhythmicity promotes metabolic abnormalities of the host (Altaha, 2022; Heddes *et al*., 2022). In addition, diet is a prominent modulator of gut microbial composition and was recently linked to microbial rhythmicity. Robust fecal microbial rhythms were detected under chow diet, whereas suppressed rhythmicity was observed in mice kept under metabolic challenging diets like high fat diet (HFD) (Leone et al., 2015; Thaiss et al., 2014; Zarrinpar et al., 2014). However, results obtained from mice kept under highly controlled high-fat purified diets were compared with control mice exposed to unrefined, non-standardized chow diets. Importantly, purified diets and unrefined chow differ dramatically in nutritional composition, specifically the content and type of fiber (Morrison et al., 2020; P.J, 2008; Pellizzon and Ricci, 2018). Purified diets contain insoluble fiber (cellulose), whereas chow diet consist both insoluble and soluble fibers, the latter being fermented by gut microbiota, which leads to production of metabolites important for metabolism (e.g. SCFAs) (Pellizzon and Ricci, 2018). Due to these compositional differences in purified diets and chow diets, it is unclear whether the observed suppression in gut microbial rhythmicity under HFD are driven by differences in dietary fat load or fiber content. Interestingly, we previously identified the intestinal clock as a major driver of fiber-fermenting microbiota, including well-known SCFAs-producer involved in metabolic homeostasis (Heddes *et al*., 2022; Lagkouvardos et al., 2019). This led us to the hypothesis that the intestinal clock mediates host metabolism by regulating fiber-fermenting microbiota.

Here we provide the first demonstration that microbial rhythmicity critically depends on the intestinal clock and on the amount of soluble fiber. Furthermore, we reveal that a functional intestinal clock is required for metabolic homeostasis of the host, particularly by regulating jejunal lipid and glucose metabolism and that this effect was additive to the effect of the intestinal clock in driving rhythmicity of fiber fermenting taxa. Together these results highlighting the physiological relevance of a functional intestinal clock in metabolic health.

## Results

### Intestinal clock-controlled microbial rhythmicity is fiber dependent and affects body weight

We recently demonstrated the importance of the intestinal clock in regulating rhythmicity of microbiota composition and function (Heddes *et al*., 2022). Specifically the highly abundant phylum Firmicutes, a well-known fiber fermenter, is controlled by the intestinal clock (Heddes *et al*., 2022) and the abundance of Firmicutes is highly sensitive to dietary alterations. Recent literature reported dampening of microbial rhythms upon high-fat diet (HFD) experimental dietary interventions in comparison to high amplitude microbial rhythms in mice under chow diet (Leone *et al*., 2015; Zarrinpar *et al*., 2014). However, these experimental diets are not only enriched in fat content, but additionally depleted of fermentable fiber, whereas the chow diet used as control contains high amounts of fermentable fibers (P.J, 2008; Pellizzon and Ricci, 2018). Although it was concluded that microbial rhythmicity depends on dietary fat content, we aimed to test whether dietary fiber could be an additional factor relevant for microbial rhythmicity. To address this, we exposed the same cohort of control mice (*Bmal1^IECfl/fl^*) and mice genetically lacking the essential clock gene *Bmal1* in the intestine (*Bmal1^IEC-/-^*) to a chow diet (23% fiber) followed by control diet (CD, low fermentable fiber, 9% fiber) and high-fiber supplemented CD (CFi, 26% fiber) dietary intervention (Fig. 1A, B). All of these diets have a similar amount of fat (∼5%), but differ in their percentage of digestible fiber (Fig. 1B). After two weeks of each diet intervention 24h fecal profiles were collected (Fig. 1A). 16S rRNA sequencing analysis showed clear differences in microbiota diversity between the diets and the genotypes (Chow p=0.065, CD p=0.001, CFi p=0.001) illustrated by beta-diversity and Generalized UniFrac distances (GUniFrac) quantification to CT1 (Fig. 1C, D). Moreover, reduced alpha-diversity (species diversity) in both genotypes depended on the diets and mirrors the amount of fermentable fiber in the different diets (Fig. 1E). Interestingly, highest amounts of circadian rhythmicity on taxonomic and zOTU level were found in control mice fed chow, followed by CFi, and lowest in low-fiber CD conditions (Fig. 1F- H), suggesting that microbial rhythmicity is indeed fiber-dependent. Of note, the dominant phases of oscillating zOTUs differed between chow (CT10 and CT22) and CFi (CT14-16) (Fig. 1H). In confirmation with our previous study (Heddes *et al*., 2022) intestinal clock deficiency caused a 40% reduction in the amount of oscillating zOTUs (and other taxonomic levels) under chow conditions (Fig. 1G-J). Importantly, although similar genotype differences were observed under fiber-rich diets, genotype differences in microbial rhythmicity were diminished under CD (Fig. 1G, I). Particularly in the absence of fermentable fiber, a substantial suppression of microbiota rhythmicity in control mice was reported, which led to a similar low amount of microbial rhythmicity in both genotypes (zOTUs: 15% control, 7%, *Bmal1^IEC-/-^*) (Fig. 1F-J). For example, highly abundant phyla, including Firmicutes and Bacteroidetes, showed circadian rhythms in control mice under Chow and CFi conditions, which was absent under CD (Suppl. Fig. 1A). Consistently, robust rhythmicity of several intestinal clock controlled families involved in fiber fermentation, including *Rumminococcaceae* and *Oscillospiraceae,* as well as several zOTUs including *Colidextibacter* and *Bacteroides* (Heddes *et al*., 2022) in control mice kept under chow as well as CFi was completely lost in CD conditions, reflecting results obtained from *Bmal1^IEC-/-^*mice (Fig. 1K, L). Prolonged fiber-depletion for the total duration of 12 weeks (Fig. 1A) dramatically augmented the reduction in microbial rhythmicity in both genotypes (zOTUs: 3%, control; 4%, *Bmal1^IEC-/-^*) illustrated on all taxonomic levels (Suppl. Fig. 1B).

**Figure 1.**
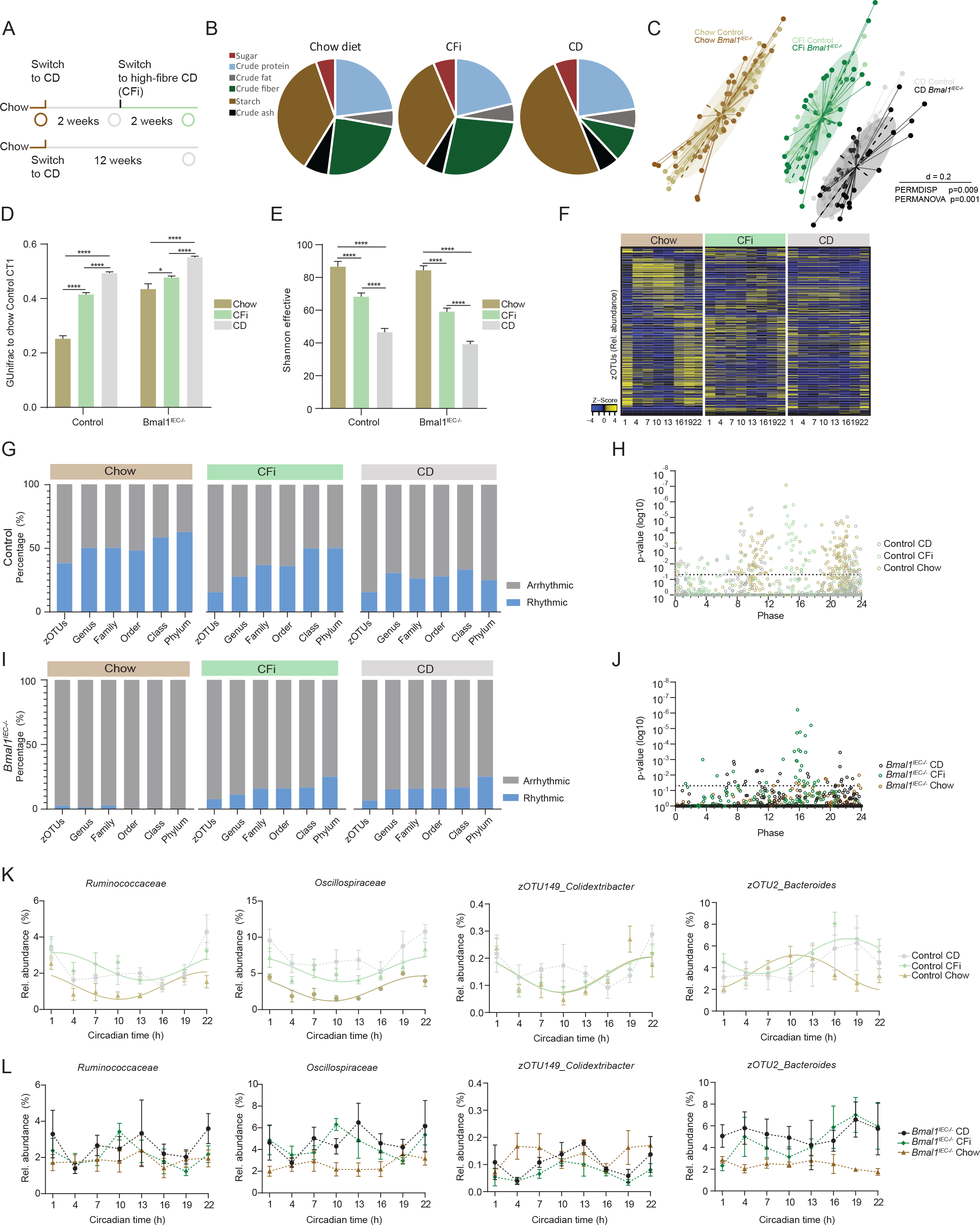
Circadian intestinal-clock controlled microbial rhythmicity is fiber dependent (A) Schematic illustration of experimental design. Circles indicate 24h fecal sampling time point. (B) Pie-charts indicating nutrient content of diets (% of total). (C) Beta-diversity illustrated by MDS plot based on generalized UniFrac distances (GUniFrac) distances of fecal microbiota. (D) GUnifrac quantification to average of chow control CT1 samples. (E) Average Alpha-diversity (Shannon). (F) Heatmap depicting the relative abundance of zOTUS in control mice. Data are ordered by the peak phase in chow. Z-score represents relative abundance where yellow means high and blue low abundance. (G) Bar-graph representing percentage of rhythmic (blue) and arrhythmic (grey) taxa in control mice identified by JTK_Cycle (Bonferroni adj. *p* value ≤ 0.05). (H) Significance (Bonferroni adj. p-value JTK) and amplitude of rhythmic and arrhythmic zOTUs and their phase distribution in control mice. (I) same as G for *Bmal1^IEC-/-^* mice and their (J) quantification of phase and p-value as described in H. (K) Circadian profiles of taxa in control mice and (L) *Bmal1^IEC-/-^* mice. Significant rhythms (cosine-wave regression, p value ≤ 0.05) are illustrated with fitted cosine-wave curves; data points connected by dotted lines indicate no significant cosine fit curves (p value > 0.05) and thus no rhythmicity. Data are represented by mean ± SEM. Significance was calculated with a one-way ANOVA within genotype. * p<0.05, **** p<0.0001.

Interestingly, the bodyweight of *Bmal1^IEC-/-^*mice under 12-week CD was significantly reduced compared to their controls (Suppl. Fig. 1C). A similar effect was already visible during two weeks of CD intervention, although it did not reach significance (Fig. 1A, Suppl. Fig. 1D). In contrast, under fiber-rich conditions, especially during the final chow intervention (week 9-11) *Bmal1^IEC-/-^* mice rapidly gained bodyweight compared to their controls (Suppl. Fig. 1D). This was validated in a new set of mice exposed to 20 weeks of chow diet, which led to a slight but significant increase in bodyweight of *Bmal1^IEC-/-^* mice (Suppl. Fig 1E). Surprisingly, the bodyweight phenotype under chow conditions was absent in a cohort kept in a different animal facility (Facility A (p=0.001), Facility B (p=0.15)) (Suppl. Fig. 1E), suggesting that the genotype effect on bodyweight might depend on the presence of specific intestinal clock-controlled microbiota. Indeed, the microbiota composition is known to substantially differ between facilities (Parker, 2018). Moreover, in the absence of microbiota, in germ-free (GF) mice, no genotype difference in bodyweight was observed, supporting the importance of intestinal clock-controlled fiber fermenting microbes on host metabolism (Suppl. Fig. 1F). It is known that the fermentation of fiber results in microbial metabolites capable of influence intestinal clock functioning (Govindarajan *et al*., 2016; Segers et al., 2019). When comparing intestinal clock gene expression under CD to chow conditions, *Per2* gene peak expression showed significant dampening in both genotypes. However, the reduction in controls was more dramatic than in *Bmal1^IEC-/-^* mice, resulting in smaller genotype differences of *Per2* expression during CD (Suppl. Fig. 1G). These results indicate that the lack of fiber might influence microbial oscillations indirectly by affecting intestinal clock function, e.g. through dampening or phase shifting. In summary, our findings highlight the importance of fermentable fiber on intestinal clock-controlled microbial oscillations and their impact on body weight. In addition, we clearly demonstrate the need to use a proper control diet to compare microbial abundances and oscillations with any purified interventional diet in metabolic studies.

### The intestinal clock regulates transcripts involved in nutrient transport and metabolism

We identified microbiota-dependent bodyweight alterations in chow fed *Bmal1^IEC-/-^* mice compared to controls, suggesting the involvement of the intestinal clock in microbial-related host metabolism (Suppl. Fig. 1). The jejunum specifically plays an important role in intestinal metabolism, by controlling nutrient absorption, digestion and regulates (microbial derived) antigen derived immune responses (Duca et al., 2021). The genotype differences in microbial rhythmicity and bodyweight gain was most pronounced in mice exposed to high fiber-containing chow diet conditions (Fig. 1). Thus, to characterize jejunal metabolic functions, transcriptional profiling was performed of jejunal (small intestine, SI) tissue sampled over the circadian cycle (24h) of control and *Bmal1^IEC-/-^* mice maintained in constant darkness kept under chow conditions (Suppl. Fig. 2A). Arrhythmic *Bmal1* and *Cry1* gene expression as well as highly suppressed *Rev-Erbα, Per2* and *Dbp* expression validated the lack of the SI clock in *Bmal1^IEC-/-^* mice, whereas control mice showed robust circadian clock gene expression (Fig. 2A, Suppl. Fig. 2B). Circadian rhythm analysis by JTK_CYCLE (Hughes et al., 2010) identified that 36% of genes underwent circadian oscillations in controls, which was reduced to 22% in *Bmal1^IEC-/-^* mice. Circadian group comparison with compareRhythms analysis (Pelikan et al., 2022) confirmed that more than 400 transcripts lost and/or changed rhythmicity in *Bmal1^IEC-/-^* mice compared to controls (Fig. 2A- B, Suppl. Fig. 2C). Although few genes remained rhythmic in *Bmal1^IEC-/-^* mice, most of them oscillated with highly suppressed amplitude, while others showed phase shifts (Fig. 2B). Of note, a small subset of transcript (41), mostly relevant for oxidative phosphorylation, gained rhythmicity in *Bmal1^IEC-/-^* mice, whereas their expression was blunted in control mice (Suppl. Fig. 2C-E, Fig. 2A). Remarkably, functional annotation of transcripts that lost rhythmicity in *Bmal1^IEC-/-^*mice, using the Panther classification system for gene ontology (GO) terms analyses, revealed overrepresentation of genes involved in metabolism. For example, intestinal clock-controlled genes were involved in pathways including ‘cellular metabolic process’, ‘lipid metabolic process’ and ‘regulation of transport’ (Fig. 2C). More specifically, GO terms involved in lipid metabolism included among others, ‘lipid homeostasis’, ‘lipid metabolic process’, ‘cellular lipid metabolic process’, ‘lipid localization’, ‘cholesterol homeostasis’ and ‘lipid transport’ were identified (Fig. 2C). Among the intestinal clock-controlled genes were the fat transporter genes, Fatty acid-binding protein 2 (*Fabp2*), Apolipoprotein B (*ApoB*) and ATP-binding cassette transporter (*Abca1*) (Fig. 2D). Moreover, genes involved in lipid digestion (*Agpat1/2, Dgat2, Acat2*) show arrhythmicity or reduced amplitudes in *Bmal1^IEC-/-^* mice compared to controls (Fig. 2D). Similarly, rhythmicity of GO terms involved in carbohydrate and protein macronutrient metabolism was lost or their amplitude was highly suppressed in *Bmal1^IEC-/-^*mice (Fig. 2C, Suppl. Fig 2F). More specifically, GO terms included ‘gluconeogenesis’, ‘hexose biosynthetic process’, ‘glucose metabolic process’, ‘monosaccharide biosynthetic process’ and ‘carbohydrate metabolic process’ (Fig. 2C). For example, the genes *Slc5a1 (Sglt1), Slc2a5 (Glut5), Slc2a2 (Glut2)*, involved in the intestinal transport of glucose, galactose and fructose to the bloodstream, showed severely reduced amplitudes or arrhythmicity in *Bmal1^IEC-/-^* mice compared to controls (Fig. 2E). Together, these data indicate that the small intestinal clock controls the circadian transcription of key metabolic-related genes involved in macronutrient absorption and transportation, particularly for carbohydrate and lipid metabolism.

**Figure 2.**
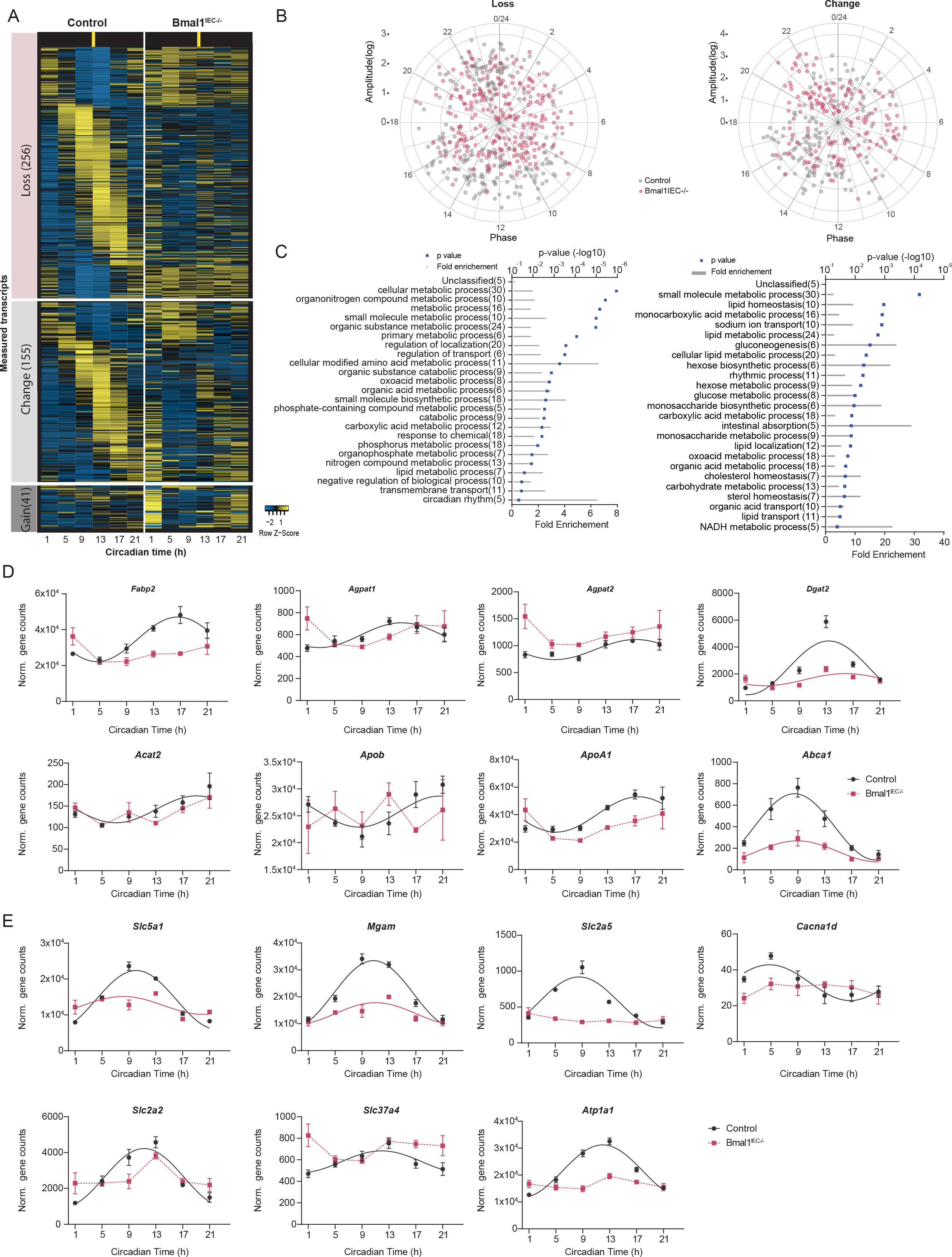
The intestinal-clock regulates the rhythmic lipid and carbohydrate transcriptome (A) Heatmap depicting altered transcripts detected by RNA-seq between control and *Bmal1^IEC-/-^*mice. Each gene (row) is organized by the phase of control mice. (B) Polarplot depicting phase and amplitude of genes loosing rhythmicity (left) and changed rhythmicity (right) in *Bmal1^IEC-/-^*mice compared to controls. (C) Pathway enrichment analyses using genes that lost rhythmicity (left) or changed rhythmicity (right) in *Bmal1^IEC-/-^*mice compared to controls. Circadian profiles of lipid (D) and carbohydrate (E) related metabolic genes measured with RNA-seq, that changed or lost rhythmicity in *Bmal1^IEC-/-^* mice. Significant rhythms (cosine-wave regression, p value ≤ 0.05) are illustrated with fitted cosine-wave curves; data points connected by dotted lines indicate no significant cosine fit curves (p value > 0.05) and thus no rhythmicity. Data are represented by mean ± SEM.

### Intestinal clock dysfunction promotes metabolic abnormalities through the cecal microbiome and metabolome involved in carbohydrate and lipid metabolism

In accordance to our previous results obtained from fecal samples (Heddes *et al*., 2022), the intestinal clock controls the rhythmicity of cecal microbiota composition (Fig. 3A). The amount of rhythmic zOTUs is highly suppressed in *Bmal1^IEC-/-^* mice and predominantly includes taxa from the highly abundant families *Muribaculaceae*, *Rumminococcaceae* and *Lachnospiraceae* (Fig. 3A, Suppl. Fig. 3A) involved in host metabolism (Barouei et al., 2017; Lagkouvardos *et al*., 2019). Transcriptomics identified dampened rhythmicity of Nuclear Factor, Interleukin 3 Regulated (*Nfil3)*, known to regulate microbial controlled lipid metabolism (Suppl. Fig. 2G) (Wang et al., 2017). Thus to further investigate intestinal clock controlled host and microbial derived metabolites we performed untargeted metabolomics (TripleTOF) of cecal content collected every 4h over a 24h period of control and *Bmal1^IEC-/-^* mice. Significant alterations in metabolites involved in taurine-, tryptophan-, vitamin B6 metabolism as well as bile acid (BA) biosynthesis and lysine-, valine/leucine/isoleucine degradation were measured in *Bmal1^IEC-/-^* mice compared to control mice (Fig. 3B). These results are in accordance to results obtained from fecal samples (Heddes *et al*., 2022). Although PCA analysis revealed no clear genotype separation, clustering within all timepoints was observed in control while *Bmal1^IEC-/-^* samples extended across multiple control cluster (e.g. CT1, CT5) (Fig. 3C). Accordingly, in the control group from all identified metabolites visualized in a heatmap (Fig. 3D), 3686 metabolites were identified to be rhythmic by JTK_CYCLE. However, in the absence of a functional intestinal clock, more than 30% of these metabolites lost rhythmicity in *Bmal1^IEC-/-^* mice. Enrichment analysis of annotated metabolites further revealed that intestinal clock-controlled metabolites are involved in amino acid metabolism (valine, leucine and isoleucine biosynthesis, pantothenate and CoA biosynthesis, aminoacyl-tRNA biosynthesis), nucleotide metabolism (pyrimidine metabolism), phenylalanine metabolism, metabolism of cofactors and vitamins (riboflavin metabolism, nicotinate and nicotinamide metabolism) and lipid metabolism (biosynthesis of unsaturated fatty acids, sphingolipid metabolism) (Fig. 3E). Interestingly, the intestinal clock predominantly controls metabolites related to carbohydrate metabolism, including galactose-, amino sugar-, nucleotide sugar-, and starch and sucrose metabolism (Fig. 3E, 3F). Particularly, mono- and disaccharides as well as hexose-phosphate lost rhythmicity in *Bmal1^IEC-/-^* mice as well as long-chain fatty acids (Fig. 3F). These results indicate the importance of intestinal clock controlled host and microbial metabolites for the host’s metabolism.

**Figure 3.**
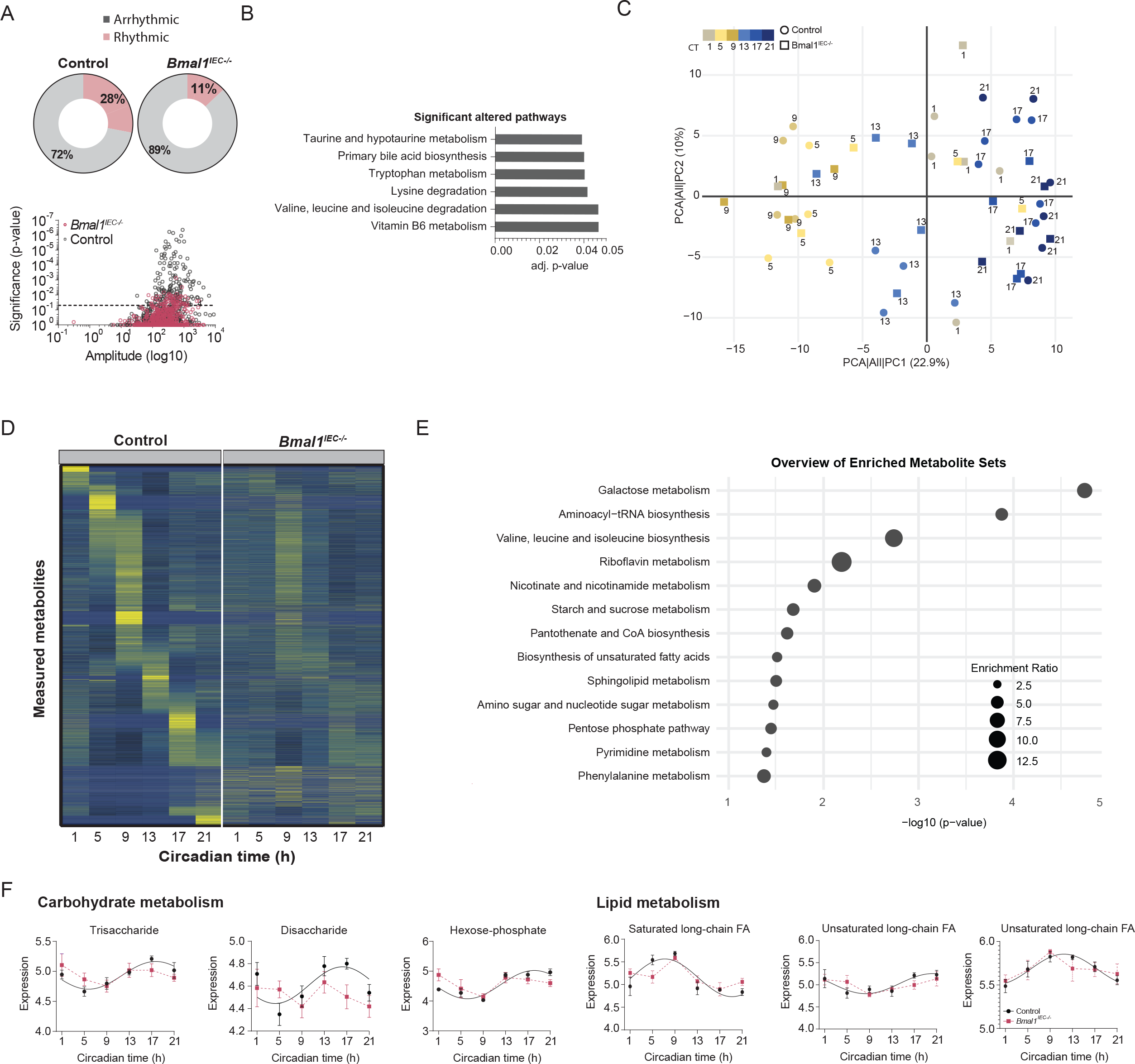
Intestinal clock control of host and microbial metabolome is relevant for carbohydrate and lipid metabolism. (A) Pie-charts depicting amount of rhythmic and arrhythmic percentage of zOTUs in the cecal microbiome, detected by JTK_cycle (Bonferroni adj. *p* value ≤ 0.05) (top). Significance and amplitude of rhythmic and arrhythmic zOTUs. (B) KEGG-pathway analyses of differentially expressed metabolites identified by untargeted metabolomics in *Bmal1^IEC-/-^* mice compared to control. (C) Principal component analyses of metabolites measured in HILIC negative ionization mode. Colors depict time and shape depict genotype. (D) Heatmap depicting all metabolites measured by untargeted metabolomics, organized by phase of control mice. (E) KEGG-enrichment analyses of metabolites loosing rhythmicity in *Bmal1^IEC-/-^* mice. (F) Carbohydrate and lipid related metabolites loosing rhythmicity in *Bmal1^IEC-/-^*mice. Significant rhythms (cosine-wave regression, p value ≤ 0.05) are illustrated with fitted cosine-wave curves; data points connected by dotted lines indicate no significant cosine fit curves (p value > 0.05) and thus no rhythmicity.

To directly test whether intestinal clock-controlled metabolites and microbiota are capable of influencing host metabolism, germ-free wild type mice (harboring a functional host clock) were gavaged with cecal content obtained from *Bmal1^IEC-/-^* and rhythmic control mice. Indeed, from 4 weeks on after the transfer wild type recipients colonized with *Bmal1^IEC-/-^* microbiota significantly gained bodyweight compared to wild types receiving wild type microbiota (Fig. 4A, B). Accordingly, dissected fat pad mass (eWAT and iWAT) was significantly enhanced, which was further confirmed by NMR showing an overall increased fat mass and decreased lean mass (Fig. 4C, Suppl. Fig. 3B). Of note, the expression of metabolic genes at ZT13 remained comparable between recipients (Suppl. Fig. 3C). Remarkably, robust oscillations on all taxonomic levels were detected in recipients 5 weeks after colonization with control microbiota (Fig. 4D). However, transfer of *Bmal1^IEC-/-^*-derived microbiota resulted in highly suppressed microbial rhythmicity, although the host clock remained functional in wild type recipients (Fig. 4D, Suppl. Fig. 3D). Notably, several taxa known to be altered in obese mouse models, including Proteobacteria and the family *Muribaculaceae* (Wang, 2020), lost rhythmicity or were suppressed in *Bmal1^IEC-/-^* recipients compared to controls (Fig. 4E, Suppl. Fig. 3E). Additionally, the abundance of *Lactobacillus,* capable of reducing bodyweight (Wang et al., 2020), was significantly increased in control recipients, while several zOTUs belonging to *Lachnospiracea* were overrepresented in *Bmal1^IEC-/-^* recipients (Suppl. Fig. 3E). Correlation analysis further revealed that the abundance of arrhythmic SCFA-producing taxa in *Bmal1^IEC-/-^* recipients, including *Lachnopiraceae* and several zOTUs belonging to *Clostridia*, positively correlated with bodyweight gain (Fig. 4E, F). A slight increase in cecal SCFA levels (p=0.06) was found at ZT13 in *Bmal1^IEC-/-^*recipients e.g. butyrate and acetate, although they did not reach significance. This increased microbial energy harvest might have contributed to bodyweight differences between genotypes (Fig. 4G). Moreover, several BAs, including the primary BAs cholic acid and β-Muricholic acid involved in fatty acid metabolism, were altered between recipients (Fig. 4H). In summary, these results highlight the importance of a functional intestinal clock in regulating microbiota and metabolite production thereby maintaining metabolic homeostasis.

**Figure 4.**
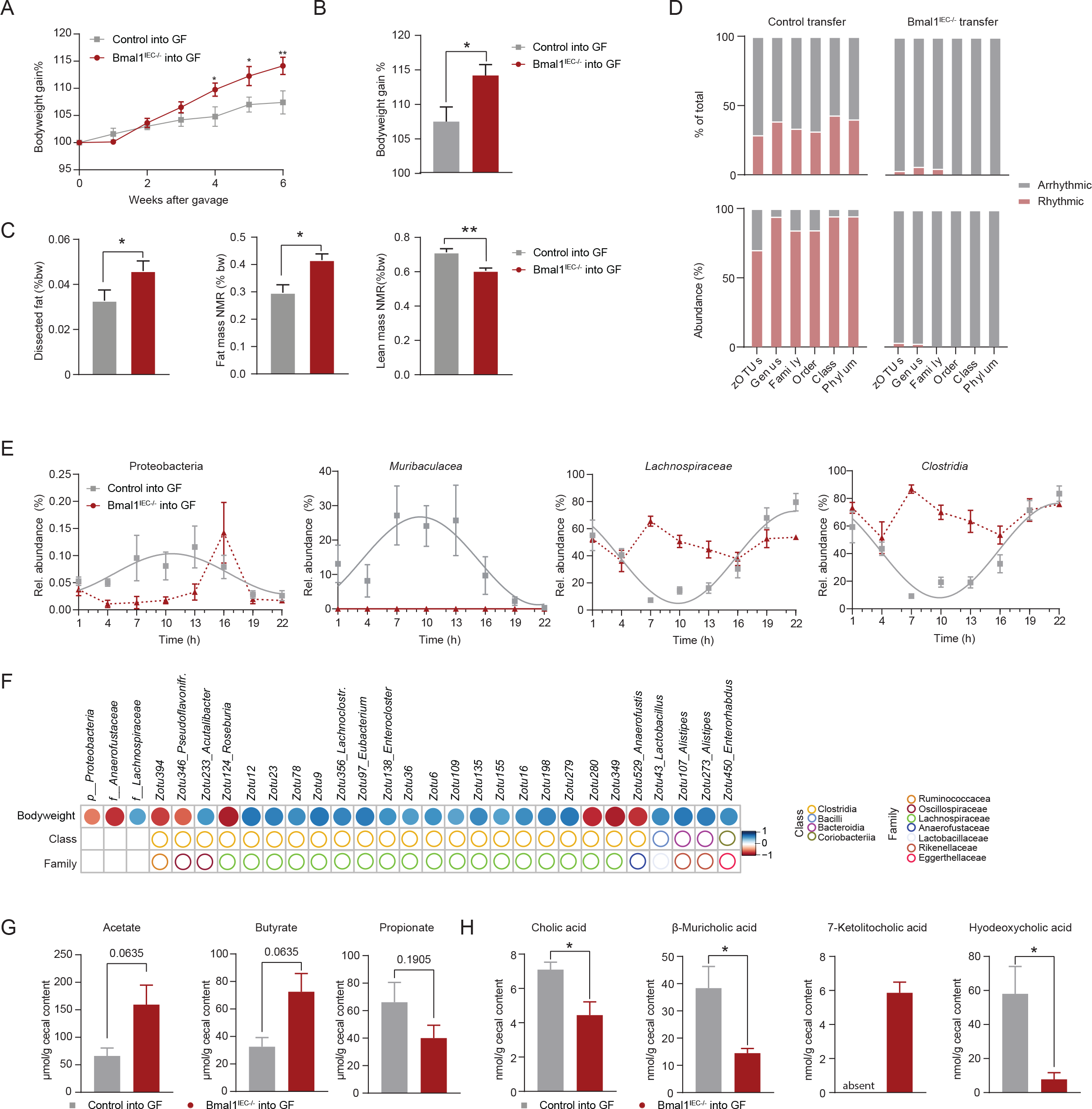
Intestinal clock dysfunction promotes metabolic abnormalities through the cecal microbiome and metabolome (A) Bodyweight gain of recipient mice after gavage with either Control or *Bmal1^IEC-/-^* cecal content. (B) Quantification of bodyweight gain at week six after gavage. (C) Fat mass measured by dissection (eWAT and iWAT) and fat and lean-mass measured by NMR, normalized to bodyweight. (D) Bar-graph representing percentage of rhythmic (red) and arrhythmic (grey) taxa in recipient mice identified by JTK_Cycle (Bonferroni adj. *p* value ≤ 0.05) (top) and their relative abundance (bottom). (E) Profiles of taxa loosing rhythmicity in *Bmal1^IEC-/-^* recipients. (F) Spearman correlation (*p* value ≤ 0.05 and R ≤ − 0.5; red or R ≥ 0.5; blue) of taxa with bodyweight. (G) SCFA and (H) BA concentrations in cecal content of control and *Bmal1^IEC-/-^* mice. Significant rhythms (cosine-wave regression, p value ≤ 0.05) are illustrated with fitted cosine-wave curves; data points connected by dotted lines indicate no significant cosine fit curves (p value > 0.05) and thus no rhythmicity. Data are represented by mean ± SEM. Significance was calculated with two-sided Mann-Whitney U test and one-way ANVOA. * p<0.05, ** p<0.01.

### Intestinal clock dysfunction promotes bodyweight gain during westernized diet feeding but not high-fat diet alone

According to our transcriptomic and metabolomics data we aim to investigate the physiological relevance of the intestinal clock for carbohydrate and lipid metabolism independent of its role on driving microbial rhythmicity. Therefore, we compared the metabolic phenotype of mice lacking intestinal clock function and controls exposed to fiber depleted high-fat (HFD, 48% kJ palm oil) and westernized diets (WD, HFD + 30% sucrose water) for 8 weeks (Suppl. Fig. 4A, Fig. 5A). In confirmation with data observed under CD (Fig. 1), the absence of fiber in HFD and WD dramatically reduces microbial rhythmicity, resulting in a comparably low amount (<8%) of rhythmic zOTUs in both genotypes (Fig. 5B-E). In contrast to recent literature (Yu *et al*., 2021) HFD-fed *Bmal1^IEC-/-^* mice and controls exhibit similar bodyweight gain (Fig. 5F, G), which was also reflected by similar weights of dissected organs and glucose tolerance test results (Suppl. Fig. 4B-D). Since we observed a significant bodyweight differences (∼1.5g) between genotypes exposed to CD (Suppl. Fig. 1B, C), weight gain under HFD was normalized to weight gain under CD in each mouse to determine the effect of the additional fat load on bodyweight gain. HFD-fed *Bmal1^IEC-/-^* mice gained slightly more bodyweight (p=0.08) in comparison to controls, although not statistically significant (Suppl. Fig 4E). This trend was independent of their nutritional intake, because the energy intake, energy excretion and total assimilation efficiency were undistinguishable between genotypes under the same dietary intervention (Suppl. Fig. 4F). Interestingly, *Bmal1^IEC-/-^*mice significantly increased bodyweight gain in compared to controls kept under HFD combined with an additional sugar load (WD) (Fig. 5H, I). Particularly, the additional sucrose intake further increased bodyweight gain in *Bmal1^IEC-/-^* mice by ∼5% normalized to HFD, whereas bodyweight gain was unchanged in controls (Suppl. Fig. 4G). Accordingly, normalized weight of dissected organs, including cecum and pancreas, significantly differed between both genotypes under WD-feeding (Suppl. Fig. 4B). Of note, sugar supplementation raised energy intake, decreased energy secretion and consequently increased assimilation efficiency in both genotypes in comparison to HFD (Suppl. Fig. 4F). Upon WD no difference was observed in total assimilation efficiency between genotypes, although the amount of fecal production and thus total energy excretion was significantly increased in *Bmal1^IEC-/-^* mice (Suppl. Fig. 4F). Nevertheless, following an oral glucose tolerance test (OGTT) a tendential higher amount of total circulating glucose (AUC) was detected in fasted WD fed Bmal*1^IEC-/-^* mice which did not reach significance (Suppl. Fig. 4B-D).

**Figure 5.**
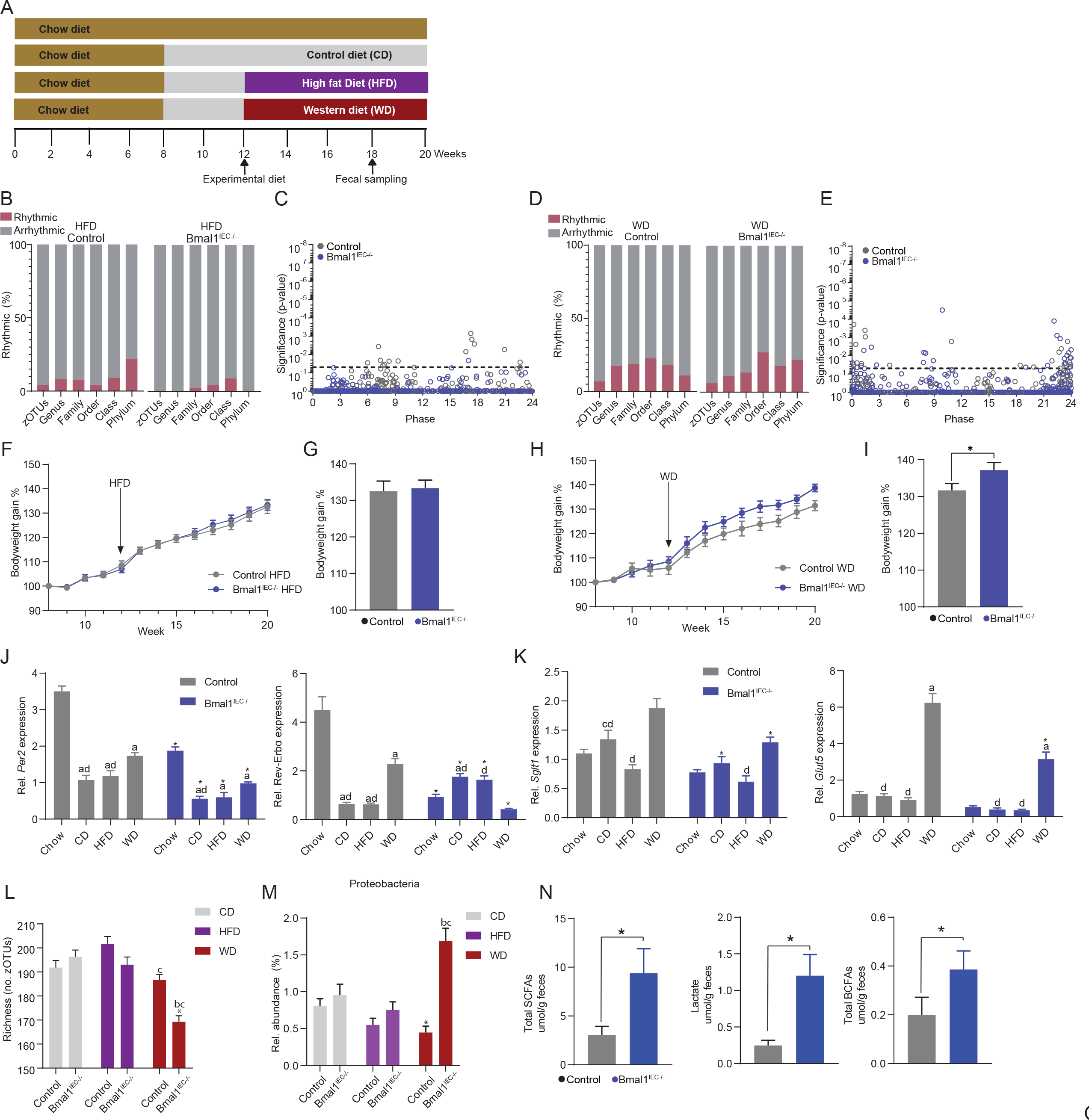
Western diet intervention promotes bodyweight gain in intestinal-clock deficient mice, independent of microbial rhythmicity. (A) Schematic diagram of experimental paradigm. (B/D) Bar-graph representing percentage of rhythmic (red) and arrhythmic (grey) fecal taxa in HFD (left) and WD (right) fed mice identified by JTK_Cycle (Bonferroni adj. *p* value ≤ 0.05). (C/E) Rhythmicity significance and phase distribution of fecal zOTUs. (F) Bodyweight gain and (G) it’s quantification of HFD fed mice and (H/I) WD fed mice. (J) Relative gene expression of *Per2, Rev-Erbα* (K) *Sglt1* and *Glut5* in the jejunum of under different dietary conditions. (L) Alpha-diversity (richness) of fecal content. (M) Relative abundance of Proteobacteria. (N) SCFA, Lactate and BCFAs concentrations in fecal content. Data are represented by mean ± SEM. Significance was calculated with two-sided Mann-Whitney U test or two-way ANOVA. In J-ML; *=significant difference between genotypes, letters mean significant difference between diets within the same genotype, with a=chow, b=CD, c=HFD, d=WD.

High caloric diet was shown to influence clock gene expression in various peripheral tissues. To investigate whether diet induced intestinal clock dysfunction caused the suppression of microbial rhythmicity in controls under HFD and WD, clock gene expression was compared at CT1 and CT13 in jejunum tissue between genotypes exposed to all dietary groups. Although, the absence of fermentable fiber in all purified diets (CD, HFD, WD) decreased clock gene expression compared to chow conditions in both genotypes (Fig. 5J, Suppl. Fig. 4H), circadian time differences between CT1 and CT13 persisted in controls, whereas no difference was detected in *Bmal1^IEC-/-^* mice (Fig. 5J, Suppl. Fig. 4H, I). Of note, in chow-fed mice increased *Rev-Erbɑ* expression was observed at CT13, whereas CT1 expression was higher in CD- and HFD-fed mice, indicating a circadian phase-shift (Suppl. Fig. 4I). These results suggest an altered, but functional intestinal circadian clock in controls and validated abolished clock function in *Bmal1^IEC-/-^* mice. Consequently, microbial arrhythmicity under fiber depleted diets was not caused by intestinal clock dysfunction in controls. Remarkably, only upon WD-feeding, reduced carbohydrate transport gene levels were detected in *Bmal1^IEC-/-^* mice compared to their controls (Fig. 5K). Notably, upregulation of genes involved in intestinal carbohydrate absorption, including *Glut5* and *Sglt1*, responsible for fructose and glucose/galactose transport respectively, were found in both genotypes under WD compared to HFD (Fig. 5K). Together these results indicate that intestinal clock dysfunction alters carbohydrate metabolism and thereby likely promotes WD-induced obesity.

Gut microbial communities are important regulators of host energy homeostasis (Wang, 2020). Although our results suggest that microbial arrhythmicity was not causal for the observed metabolic abnormalities during WD, beta-diversity differed between genotypes (Suppl. Fig 4I, J) and diets (data not shown). In WD-fed *Bmal1^IEC-/-^* mice richness was reduced, but increased abundance of the obesity-associated phylum Proteobacteria was found compared to controls (Fig. 5L, M). Moreover, the abundance of the family *Muribaculaceae* and related zOTU *Heminiphilus faecis* was reduced in controls fed WD compared to HFD and CD and this effect was even more profound in *Bmal1^IEC-/-^* mice (Suppl. Fig. 4H, I). Importantly, the abundance of *Muribaculaceae* negatively correlated with bodyweight (Suppl. Fig. 4M), supporting that in addition to circadian alterations, abundance changes of intestinal clock-controlled bacterial taxa may be linked to energy homeostasis. Indeed, microbial alterations during WD led to a significant upregulation of total SCFAs and branched-chain fatty acids (BCFAs) and lactate, whereas primary, conjugated and secondary bile-acids (BAs) show significantly reduced fecal concentration in *Bmal1^IEC-/-^* mice (Fig. 5N, Suppl. Fig. 4N, O), supporting an impact of the intestinal clock in WD-induced obesity by controlling microbial functioning.

Taken together, intestinal clock dysfunction promotes diet-induced bodyweight gain due to its role in host and microbial fat and sugar metabolism. Remarkably, this effect was additive to the role of the intestinal clock in balancing the host’s metabolism by driving rhythmicity of fiber-dependent microbiota.

## Discussion

Westernized lifestyle can result in overconsumption of dietary sugar and fat, and thereby became one of the main factors promoting metabolic disorders including obesity. Global circadian clock dysfunction was frequently linked to metabolic abnormalities, however, similar observations were made in mouse models with genetic ablation of peripheral clock function, suggesting that tissue-specific circadian clocks affect overall metabolism. For example lack of the circadian clock in the liver, adipose tissue, muscle and pancreas led to disrupted energy homeostasis, lipid and carbohydrate metabolism (Onuma *et al*., 2022; Pan et al., 2016; Paschos et al., 2012; Wada et al., 2018; Yu *et al*., 2021). Here we demonstrated that intestinal clock dysfunction promotes bodyweight gain in mice exposed to westernized diet (WD), reflected by a combination of high-fat diet (HFD) and a high sucrose solution (30%). Moreover, pancreatic weights were reduced in WD-fed *Bmal1^IEC-/-^* mice, while fasting glucose levels were slightly elevated, both associated with diabetes (Garcia et al., 2017). These results highlight the importance of the intestinal clock for carbohydrate and lipid metabolism. Mechanism likely include jejunal epithelial cell functions such as macronutrient absorption and transportation. Indeed, transcriptomic analysis revealed the importance of the jejunal clock in regulating key-metabolic genes involved in carbohydrate and lipid metabolism. In particular, the jejunal clock controls the circadian expression of hexose transporters, including *Glut5, Glut2* and *Sglt1*, which are crucial regulators for the uptake of fructose, glucose and galactose and the genes *Fabp2* and *Dgat2* previously reported to mediate intestinal fat absorption (Yu *et al*., 2021). Consistently, intestinal *Bmal1* was found to influence glucose and fat absorption by controlling diurnal *Sglt1* and *Dgat2* expression (Onuma *et al*., 2022; Yu *et al*., 2021). Sugar load was found to induce *Glut5* RNA and protein expression in the murine jejunum (Soleimani and Alborzi, 2011). This enhanced *Glut5* induction as well as enhanced *Sglt1* expression was indeed found in control mice upon WD-feeding. Interestingly, the induction of both genes was significantly suppressed in *Bmal1^IEC-/-^* mice, indicating intestinal clock regulation of jejunal specific sugar absorption. Altogether these findings demonstrate that jejunal clock dysfunction causes metabolic abnormalities under exposure to abnormal dietary glucose and fat levels. Surprisingly, HFD alone did not increase bodyweight gain in *Bmal1^IEC-/-^* mice and reduced bodyweight was observed under low fat-low fiber dietary intervention, indicating intestinal clock regulation of fat metabolism. Conversely, lack of intestinal *Bmal1* was previously reported to protect from HFD-induced obesity (Yu *et al*., 2021). Differences in the metabolic phenotype of the same mouse model between studies may be explained by different experimental conditions, such as the amount and type of fat and the feeding duration. Reduced weight gain under HFD was found when *Bmal1^IEC-/-^* mice received 60% cholesterol-rich lard fat for a duration of 10 weeks (Yu *et al*., 2021), whereas in our study 8 weeks of 48% of low-cholesterol palm oil was fed (Kubeck et al., 2016). The comparability between studies may be impacted by additional factors influencing the host’s metabolism. These include the microbial ecosystem, which is particularly relevant for dietary interventions, and was frequently reported to vary dramatically between facilities (Parker, 2018). Indeed, the bodyweight of microbiome-depleted germ-free (GF) *Bmal1^IEC-/-^*mice was undistinguishable from controls, whereas specific pathogen free (SPF) microbial-rich mice developed a significantly increased bodyweight, indicating the importance of specific intestinal clock-controlled microbiota for host metabolism.

The gut microbiota regulates energy homeostasis through multiple mechanisms, including microbial-derived metabolites. Recently we reported that the intestinal clock drives more than 50% of fecal microbial oscillations. Consistently we found that cecal microbiota composition and microbial metabolites undergo circadian rhythmicity and around half are controlled by the intestinal clock. Untargeted metabolomics further demonstrated that 30% of the circadian cecal metabolome lost rhythmicity in *Bmal1^IEC-/-^* mice, particularly host and microbial derived metabolites involved in lipid and carbohydrate metabolism. These intestinal clock-controlled pathways include galactose metabolism, biosynthesis of unsaturated fatty acids and starch and sucrose metabolism. Moreover, several bacterial species relevant for metabolic health, including Proteobacteria and families including *Muribaculacea* and *Lachnospiracea,* are under intestinal-clock control. Consequently, changes in the microbiome of intestinal clock-deficient mice likely impacted their metabolic phenotype. Dampened rhythmicity of the by microbes regulated gene *Nfil3,* regulating rhythmic transport and absorption of lipids in IECs, (Wang *et al*., 2017) was found in *Bmal1^IEC-/-^*mice, suggesting the microbiome as an important link between the intestinal circadian clock and lipid metabolism. Indeed, transfer of intestinal clock-controlled (arrhythmic) microbiota and metabolites resulted in an obesity phenotype in recipients, reflected by increased bodyweight and substantially increased fat mass. Consistently, previous studies, including our own, detected metabolic alterations in recipient mice following transfer of microbiota collected from humans and mice during circadian disruption (Altaha, 2022; Thaiss *et al*., 2014). In accordance to findings obtained from SPF *Bmal1^IEC-/-^* mice, families like *Muribaculacea* and *Lachnospiraceae*, lost rhythmicity in *Bmal1^IEC-/-^* recipient. Consequently, the amount of microbial metabolites, including SCFAs and BAs, known to play an important role in carbohydrate and fatty acid metabolism, depended on the genotype of the donor. *Bmal1^IEC-/-^* recipients showed an increase in microbial energy harvest of SCFAs, while bacterial derived secondary BA capable of reducing bodyweight gain and improving glucose metabolism (Makki et al., 2023), including hyodeoxycholic acid (HDCA), were significantly reduced. These findings highlight that intestinal clock-controlled bacterial-derived metabolites can influence obesity.

In addition to the intestinal clock, we provide the first demonstration that circadian microbial rhythmicity can be modulated by dietary fiber. The intestinal clock mainly regulates microbial rhythmicity of fiber fermenting taxa (Heddes *et al*., 2022). Accordingly, a significant difference in microbial rhythmicity between control and *Bmal1^IEC-/-^* mice was only observed under fiber rich diets, e.g. chow and fiber-supplemented CD (CFi), whereas this genotype difference was completely absent under CD, HFD and WD conditions. This dramatic suppression of microbial rhythmicity in both genotypes under fiber-depleted conditions allowed us to discover that the obesogenic effects under WD conditions were directly caused by intestinal clock dysfunction and independent from its role to modulate host metabolism by driving microbial rhythmicity. Previous reports suggested that HFD suppress microbial rhythm (Leone *et al*., 2015; Zarrinpar *et al*., 2014). However, this conclusion was based on comparison of results obtained under high-caloric dietary conditions with fiber-rich chow-fed control conditions. Here we clearly demonstrate that the lack of fiber and not the increase in fat content reduces the amount of bacterial oscillations. Consequently, quantification of microbial rhythmicity under controlled experimental dietary conditions needs to be compared to a suitable purified control diet rather than chow diets, which have not been standardized between vendors (Morrison *et al*., 2020; Pellizzon and Ricci, 2018; Tuganbaev et al., 2020). Dampening of microbial rhythmicity under fiber-depleted diets in wild type mice might be a consequence of altered intestinal clock functioning. Indeed, the intestinal clock gene *Rev-Erbɑ* show a changed time-dependency at its circadian peak under fiber-depleted diets compared to fiber-rich chow diet. Accordingly, prior reports showed phase shifts in peripheral clock gene expression in Per2::Luc mice fed rapidly fermentable cellobiose fibers compared to cellulose, which is resistant to microbial fermentation (Tahara *et al*., 2018), indicating the crosstalk between peripheral clock genes and dietary fiber intake(Marques et al., 2017). Moreover, the two core intestinal clock genes *Bmal1* and *Per2* showed dampened expression compared to chow conditions, although these clock genes peak at slightly different circadian times than the time points examined. Thus, a differentiation between a phase shift and suppression of the circadian amplitude of these genes needs to be investigated in future experiments including additional circadian time points. Nevertheless, since the intestinal clock drives microbial rhythmicity, these results suggest that the suppressed gut microbial rhythmicity in control mice fed with fiber-depleted high-caloric diets (HFD and WD feeding) was likely caused by changes in intestinal clock mechanisms. Thus, stabilizing intestinal clock function during HFD feeding might restore microbial oscillations and thereby improve metabolic homeostasis. In line with this hypothesis, time-restricted feeding with HFD was shown to stabilize peripheral clock gene expression, increased microbial rhythmicity and resulted in a reduced weight gain compared to *ad libitum* fed mice (Leone *et al*., 2015; Zarrinpar *et al*., 2014). Our recent population study demonstrated that microbial rhythmicity was lost during obesity and T2D and preceded the diabetes onset, suggesting the physiological relevance of targeting human microbial rhythmicity for disease prevention (Reitmeier et al., 2020). Future human population studies are needed to validate whether targeting the intestinal clock can restore microbial rhythmicity and modulate metabolic disorders.

Taken together, this study demonstrates that the peripheral circadian clock located specifically in intestinal epithelial cells plays an important role in metabolic homeostasis. We found the first evidence that the intestinal clock regulates energy homeostasis through two distinct mechanisms, namely rhythmic intestinal nutrient transport and the microbiome, thereby influencing the host’s metabolic response to diet. Moreover, our results highlight the importance of dietary fiber for circadian regulation of the microbiome, providing a novel link between diet, microbial rhythmicity, the intestinal clock and host metabolism.

Additional information (containing supplementary information line (if any) and corresponding author line)

## Acknowledgement

The Technical University of Munich provided funding for the ZIEL Institute for Food & Health, animal facility support, technical assistance and support for 16S rRNA gene amplicon sequencing. We are thankful for Kinga Balázs providing assistance with RNA sequencing analysis. Robert Blamberg, Qu Quojing and Johanna Bruder provided assistance with animal experiments and data collection.

## Author contributions

MH and SK conceived and designed the study. MH performed mouse experiments and data analyses. YN collected rRNAseq tissue and support to rRNAseq analyses. BA contributed to germ free mouse colonization. KK, CM and MH performed targeted and untargeted metabolomics and data analyses. SK coordinated the project and supervised data analyses. SK and DH secured funding. MH and SK wrote the manuscript. All authors reviewed and revised the manuscript.

## Material and Methods

### Animal experiments

#### Ethics statement

Experiments were conducted at Technical University of Munich in accordance with Bavarian Animal Care and Use Committee (TVA ROB-55.2Vet-2532.Vet_02-18-14).

#### Mouse models

Male epithelial intestinal epithelial cell-specific Bmal1 knock-out (Bmal1fl/fl x Villin CRE/wt; referred to as *Bmal1^IEC-/-^*) mice and their control littermates (Bmal1fl/fl x Villin wt/wt; referred to as Control) on a genetic C57BL/6J background were generated as previously described (SOURCE OUR PAPER). All mice were kept in a 12h light/12h dark cycle (300 lux), with lights turned on at 5am (Zeitgeber time (ZT0) to 5pm (ZT12)). Mice were housed under specific-pathogen free (SPF) conditions unless otherwise stated, according the FELASA recommendation.

#### Long term dietary interventions

All mice used in the experiments received ad libitum standard chow diet (V1124-300, Sniff, Soest, Germany) unless otherwise indicated. For the long term diet interventions, mice received chow diet till the age of 8 weeks and were then switched to a purified control diet (y-irradiated) (CD; S5745-E902, Sniff, Soest, Germany) till the age of 12 weeks. At the age of 12 weeks, mice either remained on CD, or got a high-fat diet (HFD, 48 kJ% palm oil-based, S5745-E912, Sniff, Soest, Germany) or western diet (WD, 30% sucrose in drinking water + HFD) till the age of 20 weeks.

#### Fiber dietary intervention paradigm

Single housed *Bmal1^IEC-/-^* and control mice received ad libitum standard chow till the age of 12 weeks. After all mice were switched to a fiber-depleted purified control diet (CD; S5745-E902, Sniff, Soest, Germany) for two weeks, followed by a high-fiber CD (CFi; S5745-E918, Sniff, Soest, Germany) for two weeks. Finally, mice received two more weeks of chow diet before sacrifice. Fecal pellets were collected in the second day of darkness (DD) at the end of every second week of dietary intervention.

#### Tissue collection

All animals were sacrificed by cervical dislocation in DD, unless otherwise specified. Eyes were removed prior to dissection and tissues were collected in DNA stabilizer overnight and stored in -80 degrees until further processing. Fecal pellets and cecal content were snap frozen using dry ice.

#### Body composition measurement

Fat mass, lean mass and free fluid were measured with the use of nuclear magnetic resonance spectroscopy (NMR, Minispec, Bruker).

#### Oral glucose tolerance test (oGTT)

Baseline glucose measurements (either at ZT1 or ZT13) were performed after 6h after starvation from tail blood, with the use of a glucometer (FreeStyle, Abbott) and glucose measuring stripes (FreeStyle, Abbott). Directly afterwards, 2 mg glucose/g bodyweight (G-40% glucose concentrate, B. Braun, Melsungen, Germany) was orally administered to the mice. Blood glucose levels (mmol/L) were measured after 15, 30, 60 and 120 min of gavage. The area under the curve (AUC) were measured as follow in mg/dl:

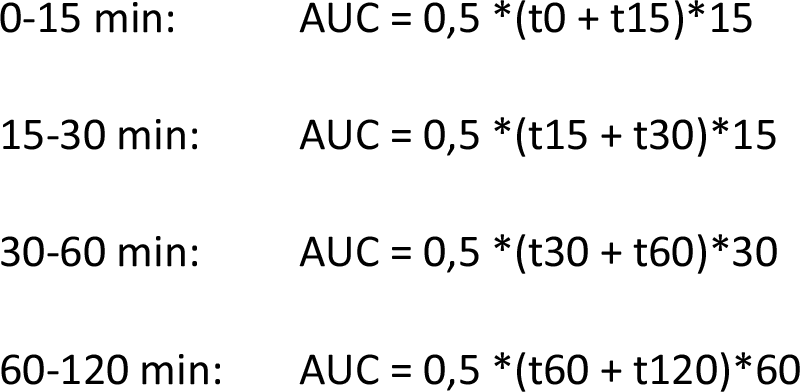
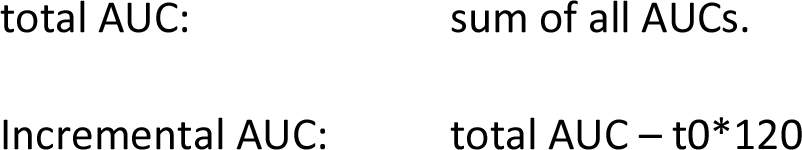

#### Microbial transfer experiments

Germ-free wild type C57BL6 were gavaged at the age of 10 weeks with a mixture cecal microbiota from either control or Bmal1IEC-/- mice cecal content diluted 1:10 in 40% glycerol. 100µl of 7x10^6^ bacteria/µl was gavaged at ZT13. After 6 weeks of the gavage, at age 16 weeks, mice were sacrificed CT/ZT13. Fecal pellets were collected every 3hours over the course of 24h day.

#### Energy assimilation

Fecal samples were collected from individual mice over 5 days and dried at 55 °C for another 5 days. Dried fecal pellets were grinded using the TissueLyserII (Qiagen, Retsch, Haan, Germany) and pressed into pellets of 1 gram (technical duplicates). Gross fecal energy content was measured using a 6400 calorimeter (Parr Instrument Company, Moline, IL, USA). Assimilation efficiency was calculated by recording the food intake and feces production over the fecal collection days as indicated in the formula below.

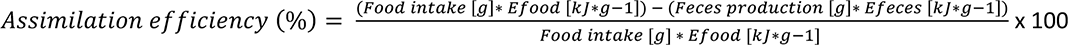

#### RNA isolation and qPCR

RNA was extracted from snap frozen tissue samples in RNAlater with the use of NucleoSpin kit according to the manufacturer’s instructions (NucleoSpin® RNA, Macherey-Nagel) and measured by NanoPhotometer®N60 (Implen). cDNA was synthesized from 1000ng RNA using cDNA synthesis kit Multiscribe RT (Thermofischer Scientific). qPCR was performed in a Light Cylcer 480 system (Roche Diagnostiscs, Mannheim, Germany) using Universal Probe Library system according to manufacturer’s instructions. For genes expression the following primers and probes were used: Brain and Muscle ARNT-Like 1 (*Bmal1*) F 5’-ATTCCAGGGGGAACCAGA-’ R 5’-GGCGATGACCCTCTTATCC-3’ probe 15, Period 2 (*Per2*) F 5’-TCCGAGTATATCGTGAAGAACG-3’ R 5’- CAGGATCTTCCCAGAAACCA-3’ probe 5, Nuclear receptor subfamily 1 group D member 1 (*Rev-Erbɑ)* F 5’- AGGAGCTGGGCCTATTCAC-3’ R 5’- CGGTTCTTCAGCACCAGAG-3’ probe 1, Cryptochrome 1 (*Cry1*) F 5’-ATCGTGCGCATTTCACATAC-3’ R 5’-TCCGCCATTGAGTTCTATGAT-3’ probe 85, Fatty acid-binding protein 2 (*Fabp2*) F 5’- ACGGAACGGAGCTCACTG-3’ R 5’-TGGATTAGTTCATTACCAGAAACCT-3’ probe 56, Sodium-Dependent Glucose Transporter 1 (*Sglt1*) F 5’-CTGGCAGGCCGAAGTATG-3’ R 5’-TTCCAATGTTACTGGCAAAGAG-3’ probe 49, Glucose transporter 2 (*Glut2*) F 5’-TTACCGACAGCCCATCCT-3’ R 5’- TGAAAAATGCTGGTTGAATAGTAAAA-3’ probe 3, Glucose transporter 5 (*Glut*5) F 5’- TGGTTGGAACCTTGGTGAATA-3’ R 5’-TGGTTGGAACCTTGGTGAATA-3’ probe 58 and RNA abundance was normalized to the housekeeping gene Elongation factor 1-alpha (*Ef1a)* F 5’- GCCAAT TTCTGGTTGGAATG-3’ R 5’-GGTGACTTTCCATCCCTTGA-3’.

#### RNA-seq

RNA quality was verified using an Agilent 2100 Bioanalyzer (Agilent) with the use of RNA 6000 Nano Kit (Agilent). Library preparation was performed with the use of Illuma TruSeq RNA library prep kit. Libraries were sequenced in a paired-end mode (2x150 bases) in the Novaseq 6000 sequencer (Illumina) with a depth of≥ 12 Million paired reads per sample.

#### Pre-processing

The quality of Next Generation Sequencing data was assessed with FastQC v0.11.5 (Andrews). Adapter content and low-quality reads were removed using Trimmomatic v0.39 (Bolger et al., 2014). Trimmed FASTQ files were then mapped against the mouse mm10 genome with the STAR v2.7.5c aligner (Dobin et al., 2013). Format conversions were performed using samtools v1.3.1 (Li et al., 2009). The featureCounts program v1.4.6 (Liao et al., 2014) was used to count reads located within an exon, do not overlap multiple features, with a threshold of MAPQ > = 4 and are not chimeric.

#### Normalization and differentially expressed genes analysis

DESeq2 version 1.22.0 (Love et al., 2014) was used to normalize the read count matrix and perform differential expression analysis. Bioconductor package ‘‘biomaRt’’ version 2.38 (Durinck et al., 2009) was used to map MGI symbols to Ensembl gene IDs. Gene ontology enrichment analyses was carried out using GOnet (qvalue<0.05) (Pomaznoy et al., 2018).

#### Circadian transcript analysis

The (differential) rhythmicity of normalized transcripts was measured through both JTK_cycle and CompareRhythms R packages. JTK_cycle (Hughes *et al*., 2010) was used through the MetaCycle R package (Wu et al., 2016) with a period of 24h and adj.p<0.05 to identify significant oscillating transcripts. CompareRhythms (Pelikan *et al*., 2022) was used to find altered rhythms between genotypes with the method DESeq2 and just_classify=FALSE. RNA-Seq datasets generated during this study are available at the GEO (NCBI) database.

#### High-Throughput 16S Ribosomal RNA (rRNA) Gene Sequencing and microbial Analysis

Genomic DNA was isolated from snap-frozen fecal pellets as previously described (Heddes *et al*., 2022) using DNA NucleoSpin gDNA columns (Machery-Nagel, No. 740230.250), followed by amplification of the V3-V4 region of the 16S rRNA gene using PCR. In short, multiplexed samples were sequenced using Illumina HiSeq in paired-end mode, with negative controls included. High-quality sequences were processed using an in-house developed NGSToolkit (Version Toolkit 3.5.2_64), which included trimming, chimera removal, merging, deduplication, clustering, and denoising steps to generate zero-radius operational taxonomic units (zOTUs). Taxonomic assignment was performed using the EZBiocloud database, and quantitative copy numbers of 16S rRNA genes were determined using artificial DNA standards. Further data analyses were performed in R with Rhea (Lagkouvardos et al., 2017).

#### Untargeted mass spectrometric measurements

Cecal samples were collected from *Bmal1^IEC-/-^* mice and their controls, every 4 hours over the course of a 24h day. Samples were directly snap frozen and stored at -80 ◦C upon metabolite extraction. The untargeted analysis was performed using a Nexera UHPLC system (Shimadzu, Duisburg, Germany) coupled to a Q-TOF mass spectrometer (TripleTOF 6600, AB Sciex, Darmstadt, Germany) as previously described (Weiss et al., 2022). Separation of the fecal samples was performed either using a UPLC BEH Amide 2.1 × 100 mm, 1.7 µm analytic column (Waters, Eschborn, Germany) with a400 µL/min flow rate or with a Kinetex XB18 2.1 x 100 mm, 1.7 µm (Phenomenex, Aschaffenburg, Germany) with a 300 µL/min flow rate. For the HILIC-separation the settings were as follows: The mobile phase was 5 mM ammonium acetate in water (eluent A) and 5 mM ammonium acetate in acetonitrile/water (95/5, v/v) (eluent B). The gradient profile was 100% B from 0 to 1.5 min, 60% B at 8 min and 20% B at 10 min to 11.5 min and 100% B at 12 to 15 min. For the reversed-phase separation eluent A was 0.1% formic acid and eluent B was 0.1% formic acid in acetonitrile. The gradient profile started with 0.2% B which was held for 0.5 min. Afterwards the concentration of eluent B was increased to 100% until 10 min which was held for 3.25 min. Afterward the column was equilibrated at starting conditions. A volume of 5 µL per sample was injected. The auto sampler was cooled to 10 °C and the column oven heated to 40 °C. Every tenth run a quality control (QC) sample which was pooled from all samples was injected. The samples were measured in a randomized order and in the Information Dependent Acquisition (IDA) mode. MS settings in the positive mode were as follows: Gas 1 55, Gas 2 65, Curtain gas 35, Temperature 500 °C, Ion Spray Voltage 5500, declustering potential 80. The mass range of the TOF MS and MS/MS scans were 50–2000 m/z and the collision energy was ramped from 15–55 V. MS settings in the negative mode were as follows: Gas 1 55, Gas 2 65, Cur 35, Temperature 500 °C, Ion Spray Voltage –4500, declustering potential –80. The mass range of the TOF MS and MS/MS scans were 50–2000 m/z and the collision energy was ramped from –15–55 V.

The “msconvert” from ProteoWizard (Adusumilli and Mallick, 2017) were used to convert raw files to mzXML (de-noised by centroid peaks). The bioconductor/R package XCMS (Smith et al., 2006)was used for data processing and feature identification. More specifically, the matched filter algorithm was used to identify peaks (full width at half maximum set to 7.5 s). Then the peaks were grouped into features using the “peak density” method(Smith *et al*., 2006). The area under the peaks was integrated to represent the abundance of features. The retention time was adjusted based on the peak groups presented in most of the samples. To annotate possible metabolites to identified features, the exact mass and MS2 fragmentation pattern of the measured features were compared to the records in HMBD (Wishart et al., 2007) and the public MS/MS database in MSDIAL (Tsugawa et al., 2015), referred to as MS1 and MS2 annotation, respectively. The QC samples were used to control and remove the potential batch effect, t-test was used to compare the features’ intensity between the groups.

The associated (un)targeted metabolomics data are available at MassIVE (MSV000091544).

JTK_cycle (Hughes *et al*., 2010) was used on all metabolites (combining positive and negative ionization mode of HILIC and RP measurements) to identify metabolite circadian oscillations. MetaboAnalyst 5.0 (Pang et al., 2021) platform was used for pathways annotation and enrichment analyses.

### Targeted metabolite analyses

#### Sample preparation for targeted metabolite analyses

Approximately 20 mg of mouse cecal content was weighed in a 2 mL bead beater tube (CKMix 2 mL, Bertin Technologies, Montigny-le-Bretonneux, France) filled with 2.8 mm ceramic beads. 1 mL of methanol-based dehydrocholic acid extraction solvent (c=1.3 µmol/L) was added as an internal standard for work-up losses. The samples were extracted with a bead beater FastPep-24TM 5G, MP Biomedicals Germany GmbH, Eschwege, Germany) supplied with a CoolPrepTM (MP Biomedicals Germany, cooled with dry ice) for 3 times each for 20 seconds of beating at a speed of 6 m/sec and followed by a 30 seconds break.

#### Targeted bile acid measurement

20 µL of isotopically labeled bile acids (ca. 7 µM each) were added to 100 µL of sample extract. Targeted bile acid measurement was performed using a QTRAP 5500 triple quadrupole mass spectrometer (Sciex, Darmstadt, Germany) coupled to an ExionLC AD (Sciex, Darmstadt, Germany) ultrahigh performance liquid chromatography system. A multiple reaction monitoring (MRM) method was used for the detection and quantification of the bile acids. An electrospray ion voltage of −4500 V and the following ion source parameters were used: curtain gas (35 psi), temperature (450 °C), gas 1 (55 psi), gas 2 (65 psi), and entrance potential (−10 V). The MS parameters and LC conditions were optimized using commercially available standards of endogenous bile acids and deuterated bile acids, for the simultaneous quantification of selected 45 analytes. A total of 27 analytes were detected in our samples. For separation of the analytes a 100 × 2.1 mm, 100 Å, 1.7 μm, Kinetex C18 column (Phenomenex, Aschaffenburg, Germany) was used. Chromatographic separation was performed with a constant flow rate of 0.4 mL/min using a mobile phase consisted of water (eluent A) and acetonitrile/water (95/5, v/v, eluent B), both containing 5 mM ammonium acetate and 0.1% formic acid. The gradient elution started with 25% B for 2 min, increased at 3.5 min to 27% B, in 2 min to 35% B, which was hold until 10 min, increased in 1 min to 43% B, held for 1 min, increased in 2 min to 58% B; held 3 min isocratically at 58% B, then the concentration was increased to 65% at 17.5 min, with another increase to 80% B at 18 min, following an increase at 19 min to 100% B which was hold for 1 min, at 20.5 min the column was equilibrated for 4.5 min at starting. The injection volume for all samples was 1 μL, the column oven temperature was set to 40 °C, and the auto-sampler was kept at 15 °C. Data acquisition and instrumental control were performed with Analyst 1.7 software (Sciex, Darmstadt, Germany) (Reiter et al., 2021).

#### Targeted short-chain fatty acid measurement

The 3-NPH method was used for the quantitation of SCFAs (Han, Lin et al. 2015). Briefly, 40 µL of the fecal extract and 15 µL of isotopically labeled standards (ca 50 µM) were mixed with 20 µL 120 mM EDC HCl-6% pyridine-solution and 20 µL of 200 mM 3-NPH HCL solution. After 30 min at 40°C and shaking at 1000 rpm using an Eppendorf Thermomix (Eppendorf, Hamburg, Germany), 900 µL acetonitrile/water (50/50, v/v) was added. After centrifugation at 13000 U/min for 2 min the clear supernatant was used for analysis. The same system as described above was used. The electrospray voltage was set to -4500 V, curtain gas to 35 psi, ion source gas 1 to 55, ion source gas 2 to 65 and the temperature to 500°C. The MRM-parameters were optimized using commercially available standards for the SCFAs. The chromatographic separation was performed on a 100 × 2.1 mm, 100 Å, 1.7 μm, Kinetex C18 column (Phenomenex, Aschaffenburg, Germany) column with 0.1% formic acid (eluent A) and 0.1% formic acid in acetonitrile (eluent B) as elution solvents. An injection volume of 1 µL and a flow rate of 0.4 mL/min was used. The gradient elution started at 23% B which was held for 3 min, afterward the concentration was increased to 30% B at 4 min, with another increase to 40%B at 6.5 min, at 7 min 100% B was used which was hold for 1 min, at 8.5 min the column was equilibrated at starting conditions. The column oven was set to 40°C and the auto sampler to 15°C. Data acquisition and instrumental control were performed with Analyst 1.7 software (Sciex, Darmstadt, Germany). Data integration and concentration calculations were done with the use of MultiQuant 3.0.3 Software (AB Sciex) as previously described (Heddes *et al*., 2022).

### Statistical Analyses

Statistical analyses were performed with the use of R and GraphPad Prism (GraphPad Software V8.0), JTK_cycle v3.1. R (25) and R. Rhythmicity analyses and differential rhythm analyses are done with the use of JTK_cycle and CompareRhythm R scripts (Hughes *et al*., 2010; Pelikan *et al*., 2022). In addition, circadian phase was calculated with the use of cosine-wave equation: y=baseline+(amplitude·cos (2·π· ((x- [phase shift)/24))), with a fixed 24-h period. Connected straight lines in of individual 24h period graphs within the figures indicate significant rhythmicity based on cosine analyses whereas dashed lines indicate non-significant cosine fit. Analysis between two groups was performed using the non-parametric Mann-Whitney test. A p value ≤0.05 was assumed as statistically significant. Rhea pipeline (Lagkouvardos *et al*., 2017) was used for microbial diversity calculations using generalized UniFrac v1.1. distances and illustrated with MDS. Heatmaps were generated using the heatmap.2 R script. Heatmaps were sorted based on peak phase of controls. Abundance plots and fold change plots were generated using SIAMCAT package in R using the “check.associations() function”. Polargraphs were made with the use of ggplot R package. Visualisation of zOTUs and samples trees were conducted using the online platform “evolgenius.info”. Spearman correlation and adjusted p-values between targeted metabolomics and zOTUs were calculated using the *rcor()* function in R. Correlation matrixes were visualized within the R package “corrplot” (Wei, 2021).

---

**Supplementary Figure 1.**
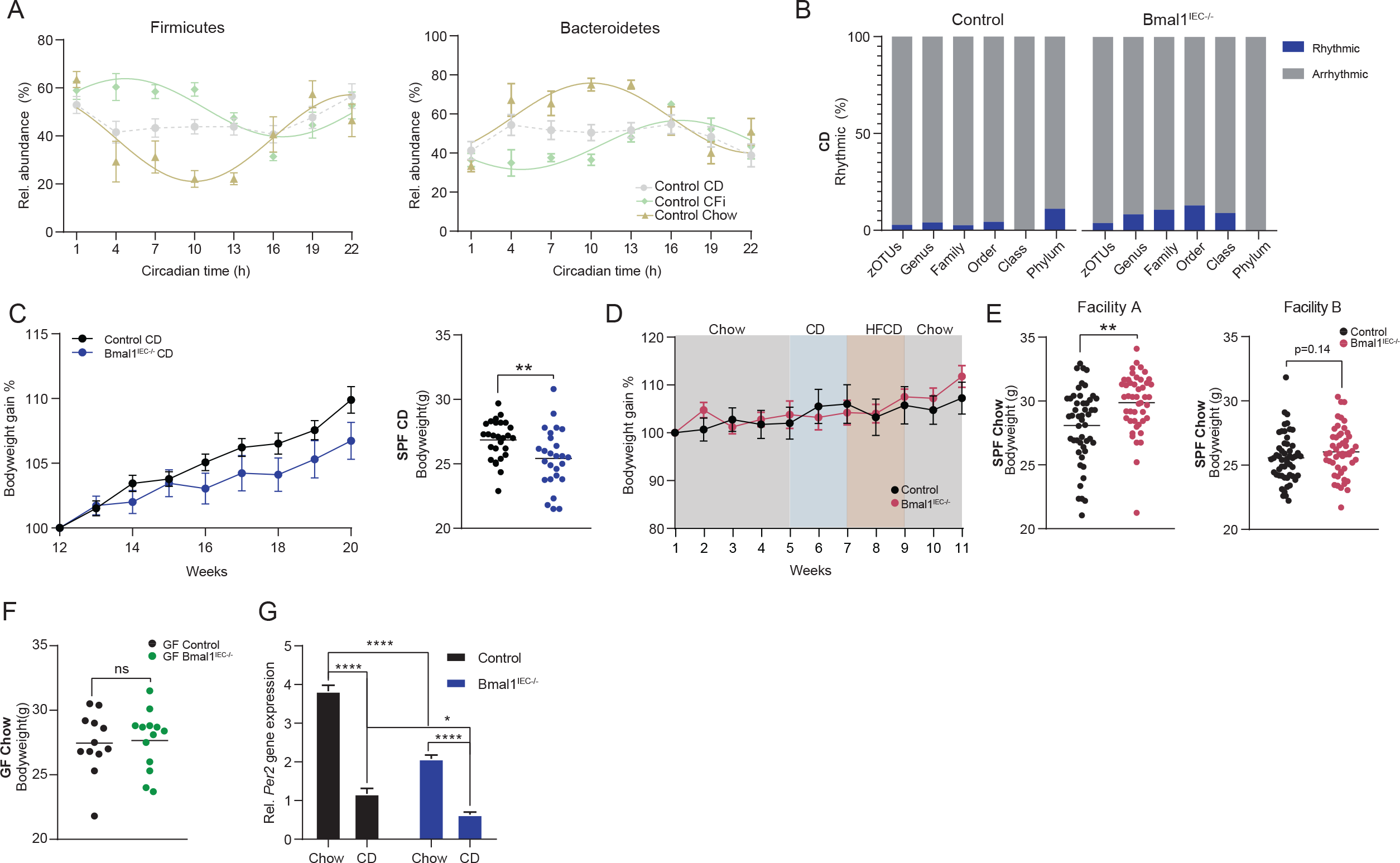
Fiber depletion decreases bodyweight in intestinal clock deficient mice. (A) Bar-graph representing percentage of rhythmic (blue) and arrhythmic (grey) taxa in CD-fed mice, identified by JTK_Cycle (Bonferroni adj. *p* value ≤ 0.05). (B) Bodyweight gain of CD-fed mice and (C) their quantification at week 20. (D) Bodyweight gain of mice undergoing altered fiber-content dietary paradigm. (E) Bodyweight of SPF chow fed mice measured in different animal facilities. (F) Bodyweight of germ-free chow fed mice. (G) Relative *Per2* gene expression in chow and CD-fed Control and *Bmal1^IEC-/-^*mice. Data are represented by mean ± SEM. Significance was calculated with two-sided Mann-Whitney U test or Two-way ANOVA. * p<0.05, ** p<0.01, **** p<0.0001.

**Supplementary Figure 2.**
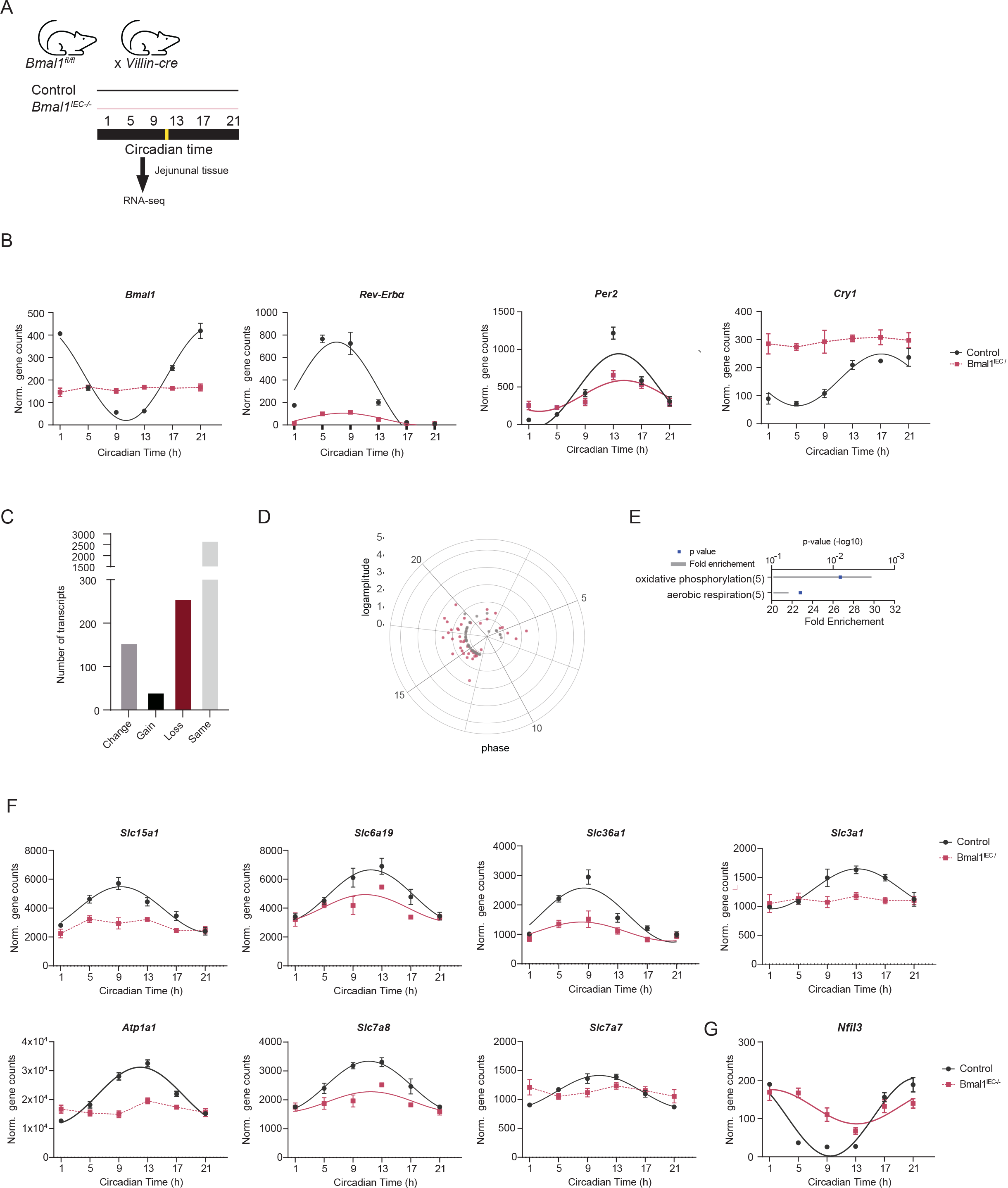
The intestinal-clock regulates the rhythmic transcriptome involved in protein metabolism (A) Schematic paradigm of RNA-seq sampled jejunal tissues (every 4-hours in constant darkness). (B) Relative clock gene expression of *Bmal1, Rev-Erbα, Per2* and *Cry1.* (C) Number of transcripts measured by RNA-seq that changed, gained or lost rhythmicity in *Bmal1^IEC-/-^* mice compared to controls. (D) Polarplot depicting transcripts that gained rhythmicity in *Bmal1^IEC-/-^*mice and their (E) KEGG-enrichment. (F) Circadian profiles of protein related metabolic genes measured with RNA-seq that gained rhythmicity in *Bmal1^IEC-/-^*mice. Significant rhythms (cosine-wave regression, p value ≤ 0.05) are illustrated with fitted cosine-wave curves; data points connected by dotted lines indicate no significant cosine fit curves (p value > 0.05) and thus no rhythmicity. Data are represented by mean ± SEM.

**Supplementary Figure 3.**
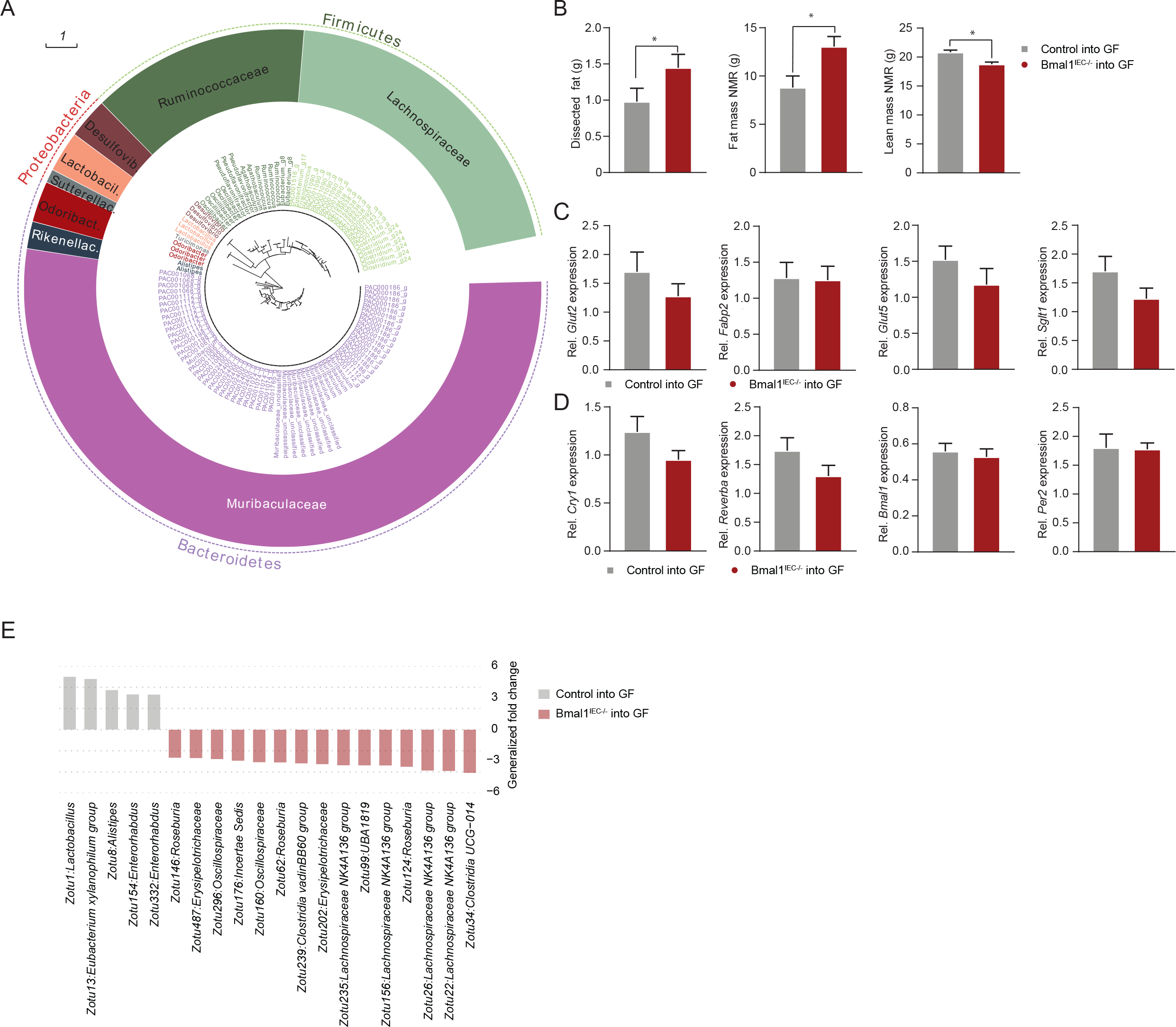
Transfer or Bmal1^IEC-/-^ derived metabolites and microbiome result in an obese phenotype. (A) Taxonomic tree of circadian cecal zOTUs under intestinal-clock control. Taxonomic ranks are from phylum (outer dashed ring), family (middle ring) to genera (inner ring). (B) dissected fat and fat and lean-mass measured by NMR in grams. (C) Relative gene expression of metabolic genes *Glut2, Fabp2, Glut5, Sglt1* and (D) clock genes *Cry1, Rev-Erba, Bmal1* and *Per2* in jejunum tissue, measured at CT13. (E) Bar plots illustrating zOTU relative abundance fold-change between *Bmal1^IEC-/-^*and Control recipients. Data are represented by mean ± SEM. Significance was calculated with two-sided Mann-Whitney U test. * p<0.05.

**Supplementary Figure 4.**
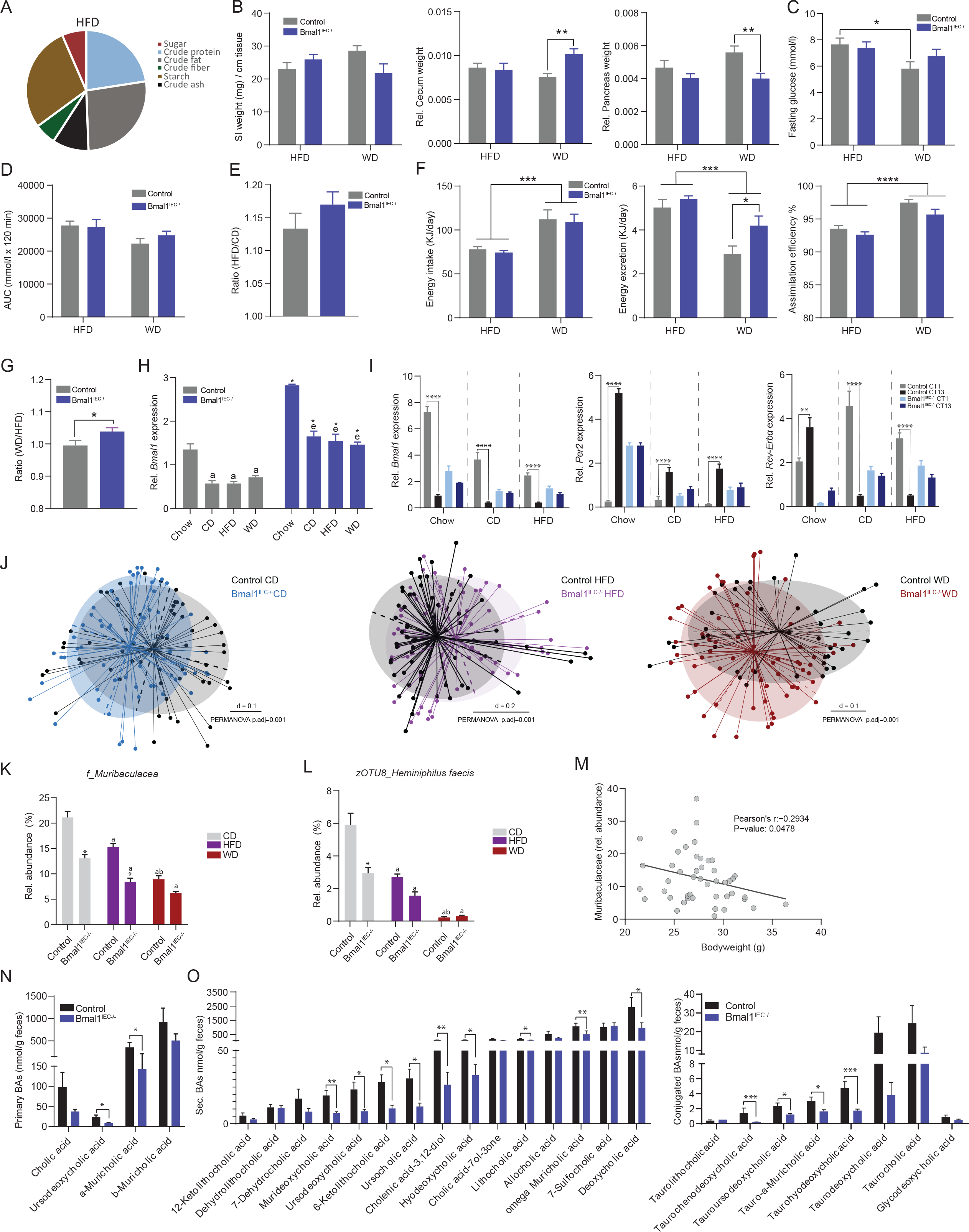
Western diet, but not high-fat diet, results in bodyweight gain in intestinal clock deficient mice. (A) Pie-chart indicating nutrient content of HFD (% of total). (B) Dissected organ weight (SI density, cecum, pancreas) relative to bodyweight after 8 weeks of HFD or WD feeding. (C) Fasting glucose and (D) total area under the curve (AUC) after glucose tolerance test. (E) Bodyweight gain of HFD fed mice, normalized to CD. (F) Energy intake, excretion and total assimilation efficiency under HFD and WD conditions. (G) bodyweight gain of WD fed mice normalized to HFD conditions. (H) Relative *Bmal1* gene expression measured at CT13. (I) Relative gene expression of Bmal1, Per2 and *Rev-Erba* at CT1 and CT13. (J) Beta-diversity illustrated by MDS plot based on generalized UniFrac distances (GUniFrac) distances of fecal microbiota. Fecal relative abundance of (K) *Muribaculaceae* and (L) *Heminiphilus faecis.* (M) Pearson’s correlation plot with bodyweight on the x-axis and relative abundance of *Muribaculacea* on the y-axis. (N) Primary and (O) secondary fecal bile-acids measured at CT13. Data are represented by mean ± SEM. Significance was calculated with two-sided Mann-Whitney U test or Two-way ANOVA. * p<0.05, ** p<0.01, *** p<0.001, **** p<0.0001. In K&L; * means significant difference between genotypes, letters mean significant difference between diet within genotype with a=CD, b=HFD, c=WD.

